# Characterizing behavioral and physiological changes in threespine stickleback (*Gasterosteus aculeatus*) with changing thermal regimes in the Canadian Pacific west coast

**DOI:** 10.1101/2025.07.15.664942

**Authors:** Angelina L. Hajji, Kelsey N. Lucas

## Abstract

Understanding the impacts of sea temperature changes on marine fishes’ physiology and behavior is critical for marine ecosystem management. Around Vancouver Island, threespine stickleback (*Gasterosteus aculeatus*) populations thrive in areas that differ by ± 10°C. To understand the acclimatory potential of stickleback, fish were exposed to cold (10°C), control (15°C), or warm (20°C) water treatments for 4 weeks. Their acclimation response was tested using the novel tank test (NTT), black-white test (BWT), measuring growth, determining critical thermal maxima (CT_max_), and measuring heat shock proteins (HSPs). We found that CT_max_ differed significantly between treatments, however, the increase in acclimation temperature was not proportional to the increase in CT_max_. In the BWT, the average number of crosses and duration that fish spent in the white zone was lower in the cold treatment than the control, while these metrics were higher in the warm treatment. A similar trend was observed in the NTT test, with warm treatment having the highest number of crosses and duration in the top third of the tank, and the cold treatment having the lowest. Surprisingly, fish in the cold treatment had a significantly greater change in weight and lowest HSP content than fish kept in the other treatments, which may be associated with energy expenditure and trade-offs for activity. Given these results, greater trade-offs in fish energy usage are anticipated in changing climates due to the acclimation limits of their bodies which would starkly affect ecosystem dynamics.

**HIGHLIGHTS:**

- Climate change is driving variability in habitats’ thermal regimes directly impacting the function and performance of animals.
- Behavioral changes (in anxiety or anti-predatory and exploratory behaviors) ensue under changing thermal environments.
- Stickleback are reaching the ends of their acclimatory capacity under current peak summer water temperatures.
- Consequences are anticipated with continued warming through trade-offs among energetically expensive processes like behavior, growth, and stress response.

## INTRODUCTION

Temperature is a dominating factor for ectothermic animals including most fishes, playing a key role in an individuals’ metabolism, enzyme functionality, physiology, and behavior (1,2). Climate is rapidly changing, biodiversity and populations are declining, and sea temperatures are rising; in Canada temperatures are rising at twice the global average (3). Climate, and particularly temperature, is the single most important determining factor for biodiversity (4) due to the integral role it plays in organism performance (5). As temperatures continue to rise, animals must compensate across all levels of biological organization (6). Thus, they may experience energetic trade-offs. Here we explore how shifting thermal tolerance may result in behavioral changes and trade-offs in energy expenditure.

Behavior is a fundamental aspect of an individual’s fitness. The behavior of an individual is integral to successfully acquiring a food resource, communication and signalling between conspecifics, reproductive success (e.g., defending a territory), and avoiding predation (7,8). Changes in behavior stem from physiological, genetic, and neurological interactions that inherently must be integrated and highly coordinated for an animal’s survival (i.e., behavior is also a physiological endpoint) (9). Thermal behavioral and physiological trade-offs are not unheard of. For instance, in mammals, the physiological mechanisms of shivering and the social behavior of huddling involve involuntary and voluntary decision-making, respectively, and are driven by (energetic) trade-offs (9). Here, trade-offs might consist of energy spent in movement to the herd or access to resources, versus energy spent on warming. Additionally, stress and anxiety-like behaviors play key roles in this decision-making process and innate trade-offs. With warming temperatures, prolonged and more frequent anxiety behavior would indicate changes to population dynamics (e.g., averseness to defend territories and find mates thereby exhibiting decreased fecundity), and decreased resilience (10–12), regardless of apparent thermal tolerances. Importantly, these behaviors have further implications towards the fish’s energy budget as more anxious individuals tend to display greater anti-predatory behavior and less foraging behavior (e.g., observed in frogs (13)). It remains unknown whether a trade-off between behavior and other physiological processes ensues in fish from shifting thermal tolerances.

In general physiological processes are highly dependent on temperature (14). Performance and function, and any associated physiological process, of an animal are expected to increase with temperature until an optimum, where after performance rapidly declines until mortality (Figure 1a; (14,15)). Enzymatic and physiological endpoints are important metrics in which functionality is lost yielding a loss in equilibrium, dysfunctionality, and potentially mortality (16). Proteins and enzymes are three-dimensional and subject to thermodynamic effects resulting in the rising segment of performance curves and denaturation effects associated with the declining segment (17). Both are contingent on temperature and are reflected in performance (14). Denaturization, the unfolding of proteins, is a time-dependent and irreversible process (i.e., longer time at higher temperatures increases denaturation risk). These processes and interactions on the molecular scale drive organism performance and functionality at higher levels of biological association.

**Figure 1.**
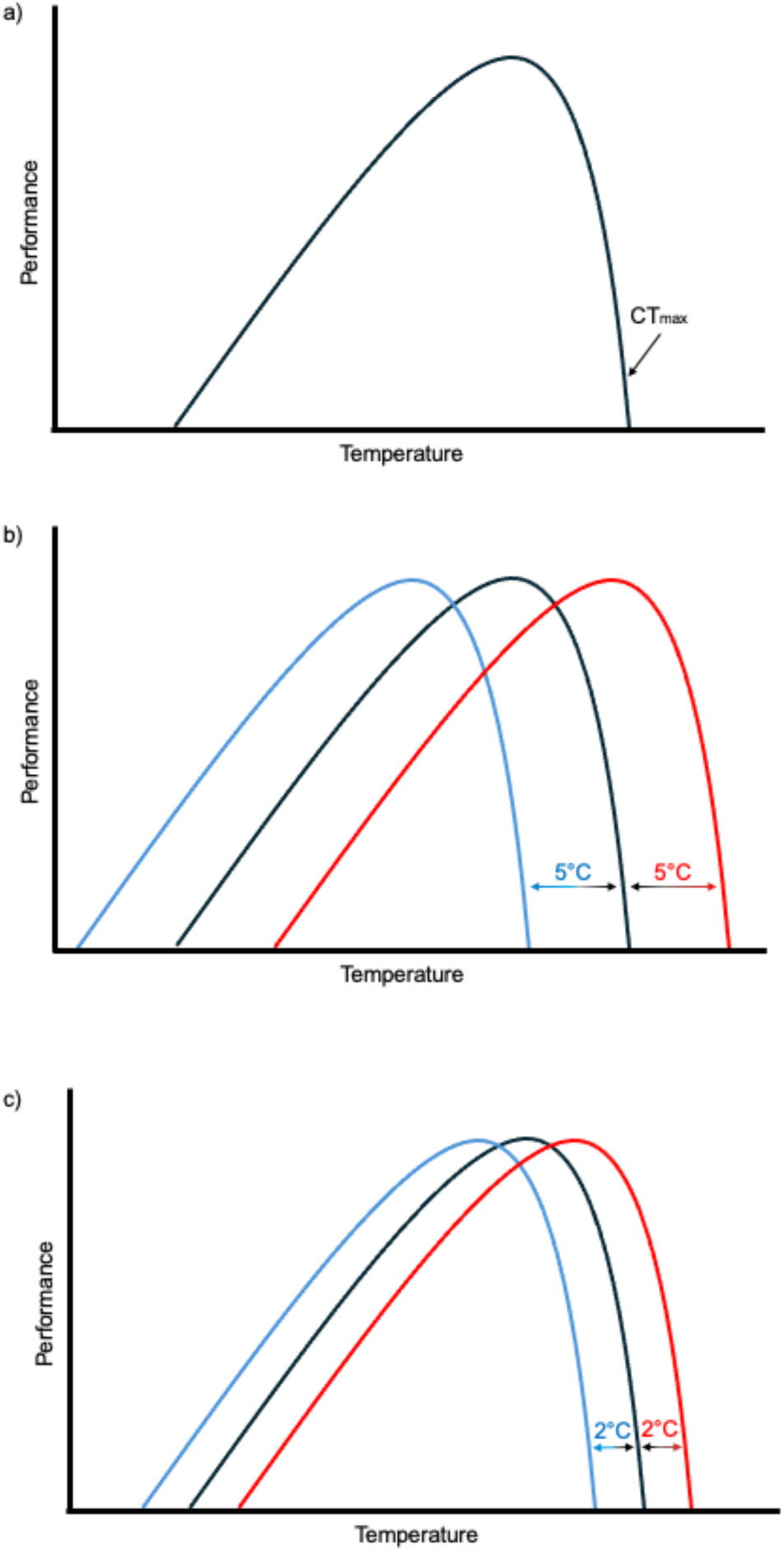
Thermal performance (function) curves with colors representing theoretical acclimation temperatures of 10°C, 15°C, and 20°C. a) a standard thermal performance curve with CT_max_ on the curve, b) example of a theoretical performance curve shift that is proportional to the ambient temperature of acclimation, and c) example of a theoretical performance curve shift that is not proportional to the ambient temperature of acclimation.

An animal’s Critical Thermal Maximum (CT_max_), the upper thermal tolerance limit, is an effective enzymatic and physiological endpoint (18,19). This point represents a loss of function or performance, and in fish, is typically taken as the temperature at which the fish loses equilibrium or the ability to maintain an upright posture (18). Death would shortly follow in natural settings due to an inability to capture prey or evade predators (20–22). CT_max_ also represents a state of physiological disorganization due to destabilization of proteins (23). In relation to the thermal curve (Figure 1a), CT_max_ occurs towards the right extreme prior to mortality. CT_max_ can vary between species, populations, and between individuals of a population (24). When exposed to different temperatures for an extended period, an individual’s CT_max_ may shift signaling a thermal acclimation response (25). Whether the change in CT_max_ is proportional to the change in acclimation temperature (e.g., both the environmental and CT_max_ increase by 5 °C) can determine the capacity and level of stress an animal at that temperature is experiencing (Figure 1b). If the change in CT_max_ is disproportional to the environmental temperature change, the shifting of the thermal curve will be reduced (Figure 1c). Functionally, the animal would be in the declines of their curve outside of their optimal temperature range (or ascending towards this) and functionality of the animal is expected to decline (Figure 1c).

Other physiological responses such as increases or decreases in the expression of heat shock proteins (HSPs) are expected to occur with changing temperatures (26). HSPs are molecular chaperones that aid protein assembly and folding, and they are activated with heat, chemical, or physical stressors and during cell generation, immuno-responses, and with cell aging (27,28). These proteins are typically mobilized towards the ends of the thermal performance curve and contribute to acclimation alongside other processes (e.g., membrane restructuring and other protein metabolism) by chaperoning proteins and allowing them to maintain performances (28). These actions under new conditions thereby allow the shifting thermal curves (29). HSPs are numbered by their molecular weight (e.g., HSP70 weighs less than HSP90). HSP70 and HSP90 expression has been characterized in fish with significant changes in expression with changing temperature, where these proteins prevent deformation of other proteins with external stress (30). The mobilization and upkeep of these proteins likely is energetically costly, which could lead to energetic trade-offs with other activities such as behavior. Characterizing these trade-offs will aid in classifying fishes’ resilience to different thermal regimes, particularly, in understanding their populations’ performance, stability, and in extreme cases, survival.

Threespine stickleback (*Gasterosteus aculeatus)* are an ideal study organism to test organismal and ecological predictions in response to temperature, as these fish are found across the globe in a variety of environments with ranges in salinity (marine, freshwater, and brackish), depth (0-100 m), temperature (4°C-20°C), and location (72°N-25°N,117°E-60°E; Froese and Pauly 2022). Due to extensive geneflow, marine populations in British Columbia are genetically conserved (genetic variation between populations mirrors within populations), and variability in phenotypes are due to acclimatory potential, i.e., phenotypic plasticity (ability to change phenotype) in response to the environment (31–34). As widespread secondary consumers they are integral to ecosystem functionality; their role as both predator and prey allows us to infer or predict trophic effects from observing stickleback responses to environmental stressors (35). Thus, the response of stickleback to different thermal regimes can provide insight to direct and indirect effects on individual and population performance in the ecosystem.

The primary purposes of this study were to 1) determine the acclimation potential and thermal capacity of stickleback at various temperatures, and 2) understand the effects of different temperatures on the behavioral responses of stickleback. We hypothesized that CT_max_ of sticklebacks will increase with increasing acclimation temperature due to their remarkable phenotypic plasticity but that there will be a limit to CT_max_ increase capabilities due to enzymatic and physiological limits. As a functional resulting trade-off for this acclimatory thermal capacity, we expected fish acclimated at higher temperatures to demonstrate decreased anxiety behaviors such as anti-predatory responses (e.g., flight response, fast starts, diving, etc.), thereby driving other energetic tradeoffs (36). As climate warms and habitats of fish move closer towards fishes’ thermal capacities, understanding changes in behavior, physiology (i.e., enzymatic), thermal tolerances and the possibility of energetic trade-offs becomes integral to determining how individuals respond to thermal stress, how energy budgets may be impacted, and what impacts on population or ecological dynamics may result.

## METHODS

### Ethics

All work was approved by the University of Calgary Life and Environmental Sciences Animal Care Committee and performed under the Animal Care Protocols AC22-0069 and AC21-0053.

### Fish Sampling and Husbandry

Adult sticklebacks (F4; ∼1 year) descending a wild marine population originally collected from sites near the Bamfield Marine Science Centre (48°49’12.69“N 125° 8’57.90“W) were bred in lab in the winter of 2022. Three families were used throughout the study. These fish were housed in the University of Calgary’s Life and Environmental Sciences Animal Resources Centre (LESARC) facility and held under standard fish lab conditions (15°C, 16D:8N, 10 ppt salinity). Fish were elastomer tagged as juveniles to provide identifiers for individuals. To initiate the experiment adult fish were randomly sorted into 9 tanks (10 L) with 5 fish per tank (N = 45). To reduce tank effects and ensure consistent family distributions between treatments, fish were distributed following a block design. After transfer between tanks, they were given one week to recover from any handling stress and to form new social hierarchies. Tanks were provisioned ∼ 2.7% g/g blood worms for the first week and 5.5% g/g for the following weeks to standardize feedings between tanks. All fish remained acclimated to a common temperature of 15°C (about the preferred temperature for stickleback (37,38)) to provide a common baseline for the individuals (see Figure 2 for experiment overview). Sticklebacks were then acclimated for 4 weeks at temperatures of 10°C (10.0 °C ± 0.29 SE), 15°C (15.1 °C ± 1.15 SE), and 20°C (20.4 °C ± 0.24 SE), with three tank replicates per temperature treatment. These temperatures were selected for their ecological relevance, to reflect current climate conditions observed from sea surface temperature trends, respectively from 1) Race Rocks in Pacific Fishery Management Area (PFMA) 20, 2) Amphitrite Point in PFMA 23, and 3) Chrome Island in PFMA 14 (see Figure 3). The sites are similar in latitude but differ in their mean and max temperatures, based on the Canadian Lightstation datasets (39).

**Figure 2.**
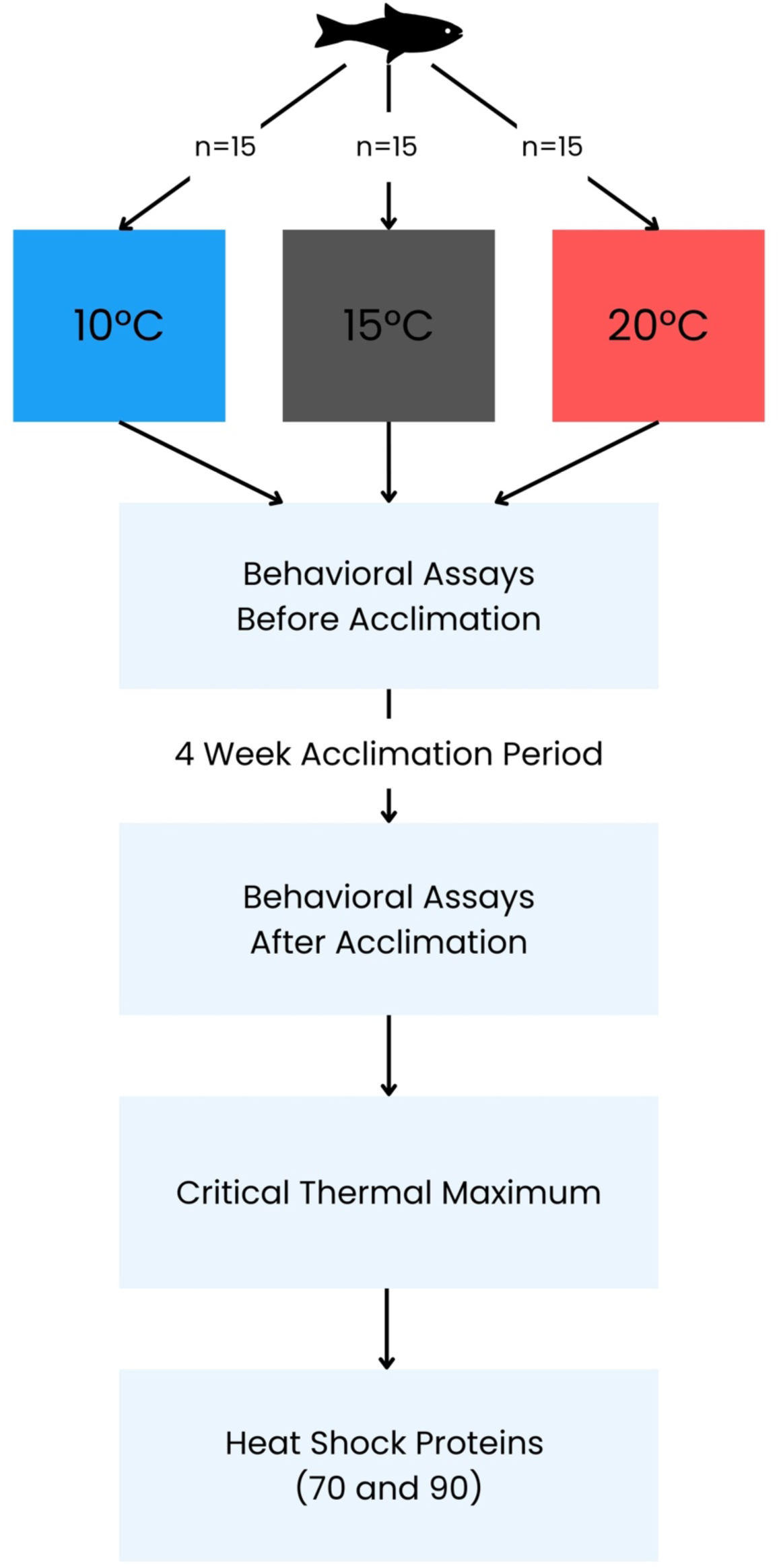
Overview of experimental design

**Figure 3.**
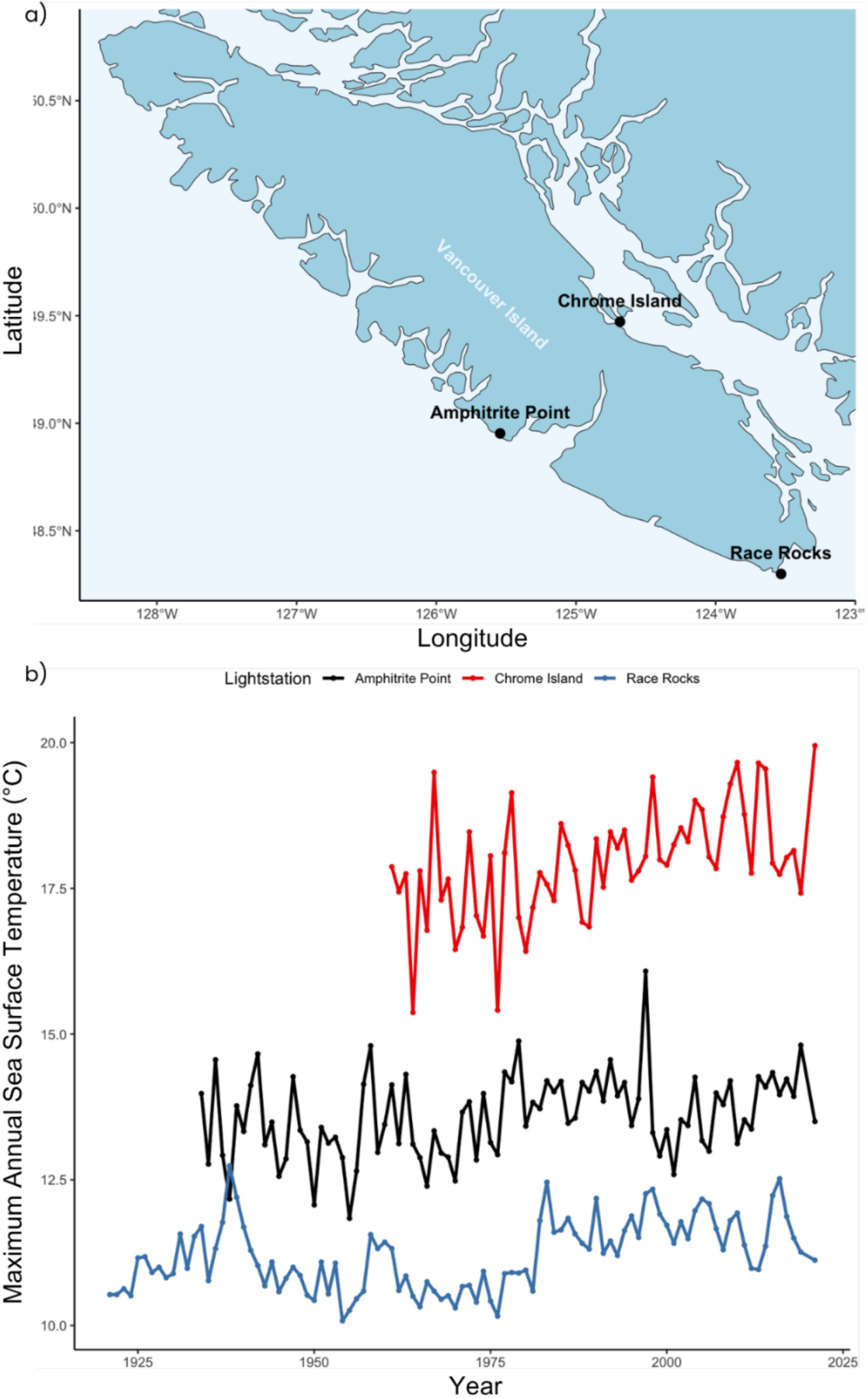
a) Government of Canada Race Rocks, Amphitrite Point, and Chrome Island Lightstation sites and b) maximum sea surface temperature (°C) over year from three sites Race Rocks (blue), Amphitrite Point (black), and Chrome Island (red), over year. Dotted lines indicate trend line from temperature points. Data taken from Government of Canada Lightstation dataset (39).

### Behavior

The responses of fish to the temperature treatments were determined through general observations and two common behavioral assays (BWT and NTT) following Norton and Gutierrez (40) (N = 45). The assays were performed before and after the 4-week acclimation period for all treatments. Briefly, for the first test single fish were transferred into a transparent tank (33.5 cm x 17 cm x 22 cm) with black and white opaque covers that divide the tank into a white zone and a black zone and recorded from above using a GoPro HERO 10 for 5 minutes (240 fps, linear mode). The behavior was manually quantified from the footage by recording the time spent in the white zone and the number of crosses made (from the black to the white zone). To avoid unintentional bias, the researcher quantified the behavior was blinded during the “after” phase, as to which treatment is which, by a secondary researcher randomizing footage filenames before measurements occurred. This was sufficient because individual fish had no defining characteristics that would allow researchers to identify their group based on appearance. For the second test, fish were transferred one at a time into a tank divided into three unmarked zones: bottom, middle, and top. They were then similarly recorded from the side for 5 minutes to measure the time spent on the top third of the tank, total distance swam, and freezing. EthoVision^®^ XT 14 (Noldus Information Technology, Wageningen, the Netherlands (41)) was used to automatically quantify these parameters and, as such, no randomization was required. For both assays, the water in the assay tanks were changed every second fish and maintained at the temperature the fish were acclimated to.

### Growth and Mortality

To minimize handling stress prior to behavior recordings, the length and weight of the fish were recorded following behavioral measurements (N = 45). This included the weight in grams and the total length of the fish in mm. These data were used to calculate the condition factor (K) using Pauly (42):

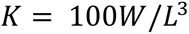

Where K is condition factor, W is the weight of the fish in grams, and L is the length of the fish in cm. We additionally used a modified the exponent of 3.04 (i.e., *K* = 100*W*/*L*^3.04^), calculated as an average of length-weight exponents from threespine stickleback data available on Fishbase.org (43) and unpublished data collected in the lab, as this value more specifically resembles their growth pattern. One mortality occurred in the warm treatment over the duration of the experiment, on day 27, which followed final behavior endpoint testing, so this fish was only excluded from CT_max_ trials.

### Critical Thermal Maximum (CT_max_)

Following the behavioral assays, the fish were fasted for 24 hours, in preparation for the CT_max_ trials, used to determine thermal tolerances. Four fish from each tank were randomly sampled for this measurement (N = 36). Briefly, to determine CT_max_, the temperature is raised constantly at the standard rate of temperature change of approximately 0.33°C min^-1^ until the fish can no longer maintain dorsoventral positioning (18,22). To ensure consistency between fish, fish were placed in 1000 mL glass jars in the experiment tank and provided a 15-minute acclimation period at the temperature of their home tank prior to the trial commencing. The thermal safety margin was calculated as the difference between CT_max_ and ambient temperature.

### Protein Expression

At the conclusion of the CT_max_ experiment, fish were euthanized. Livers were dissected from stickleback (CT_max_ individuals) and stored at −80°C. HSPs were analyzed as previously described (44,45). In brief, proteins were homogenized using sonication on liver samples in 100 uL of Tris-B and protease inhibitase solution. The samples were then centrifuged for 2 minutes and the supernatant containing the proteins were removed. To quantify the total protein concentration of samples, a bicinchoninic acid assay was performed and protein concentrations were standardized using results by adding 2X Laemmli’s buffer with *β*-mercaptoethanol (1:1). To denature proteins, samples were boiled on a heating block for 5-8 minutes and then loaded into a western blot with the first lane containing a standardized latter. Proteins separated were transferred from the wet gel to a dry membrane (semi-dry western blot). Using primary and secondary antibodies for heat shock proteins 70 (anti-HSP70 2m1) and 90 (anti-HSP90 SPC-316D), with Goat Anti-Rabbit IgG (H + L)-HRP Conjugate (1706515) as the conjugate. After imaging, to quantify data, ImageJ software (Rasband, 1997–2014) was used following Gallo-Oller (46). Briefly, image backgrounds are subtracted and the area around each band is selected and quantified for pixel intensity provided as arbitrary area values, where higher values represent greater protein expression.

### Data Analysis

To determine any significant variation in morphological parameters, CT_max_, and mortality rates among acclimation temperatures (10°C, 15°C, and 20°C), one-way ANOVAs were performed where any significant results (a = 0.05) were followed by a Tukey’s test to identify where the difference existed. Tank was initially included in each model to account for possible tank effects, but as it was not significant, it was removed from the final models through backward selection. These tests were preceded by Shapiro-Wilk’s tests to check the assumption of normality and Barlett’s tests to check the assumption of homogeneity of variance. As the assumption of normality was not met in the behavior metrics, the non-parametric Kruskal-Wallis test was performed followed by the Dunn’s test to identify where the difference existed. R (47) was used to perform all statistical tests.

## RESULTS

### Behavior

In general, researchers observed variable activity levels between temperature treatment tanks. Cold acclimated fish demonstrated lower activity levels and were often observed stationary or hiding in plants placed in the tanks. The warm tanks were comparatively more active than both the cold and control treatment tanks, and more aggressive, particularly during feedings (e.g., such as more flaring of three spines or biting of conspecifics). While routine behavior was not quantified, these general observations were reflected in results from the BWT and NTT.

In the BWT there was a significant decrease in the number of crosses from the black zone to white zone in the cold treatment compared to the control and warm treatments (Z-Score = −2.708606 and −3.792049; p = 0.0034 and 0.0001 respectively). The difference between the control and warm frequency of visits to the white zone was not significant (Z-Score = −1.083442; p = 0.1393); however, when comparing before and after within treatments, the warm treatment exhibited a significant increase (Z-Score = 3.136515; p = 0.0009). The average number of crosses observed decreased in the cold treatment tanks from 2.5 to 1.5, but this change was not significant (Z-Score = −0.776967; p = 0.2186). There was no significant differences observed in the duration of time spent in the white zone between treatment groups (Kruskal-Wallis chi-squared = 1.0056; p = 0.6048), however the average time spent in white also increased from 59.5 ± 22.7 s to 109.6 ± 18.0 s for the warm treatment (Kruskal-Wallis chi-squared = 5.6308; p = 0.3438). The opposite trend was observed for the cold treatment where the average duration spent in the white zone decreased from 133.5 ± 30.5 s to 100.7 ± 33.1 s (Z-Score = −0.973697; p = 0.1651). Figure 4 summarizes the results between treatments for the BWT behavioral assay.

**Figure 4.**
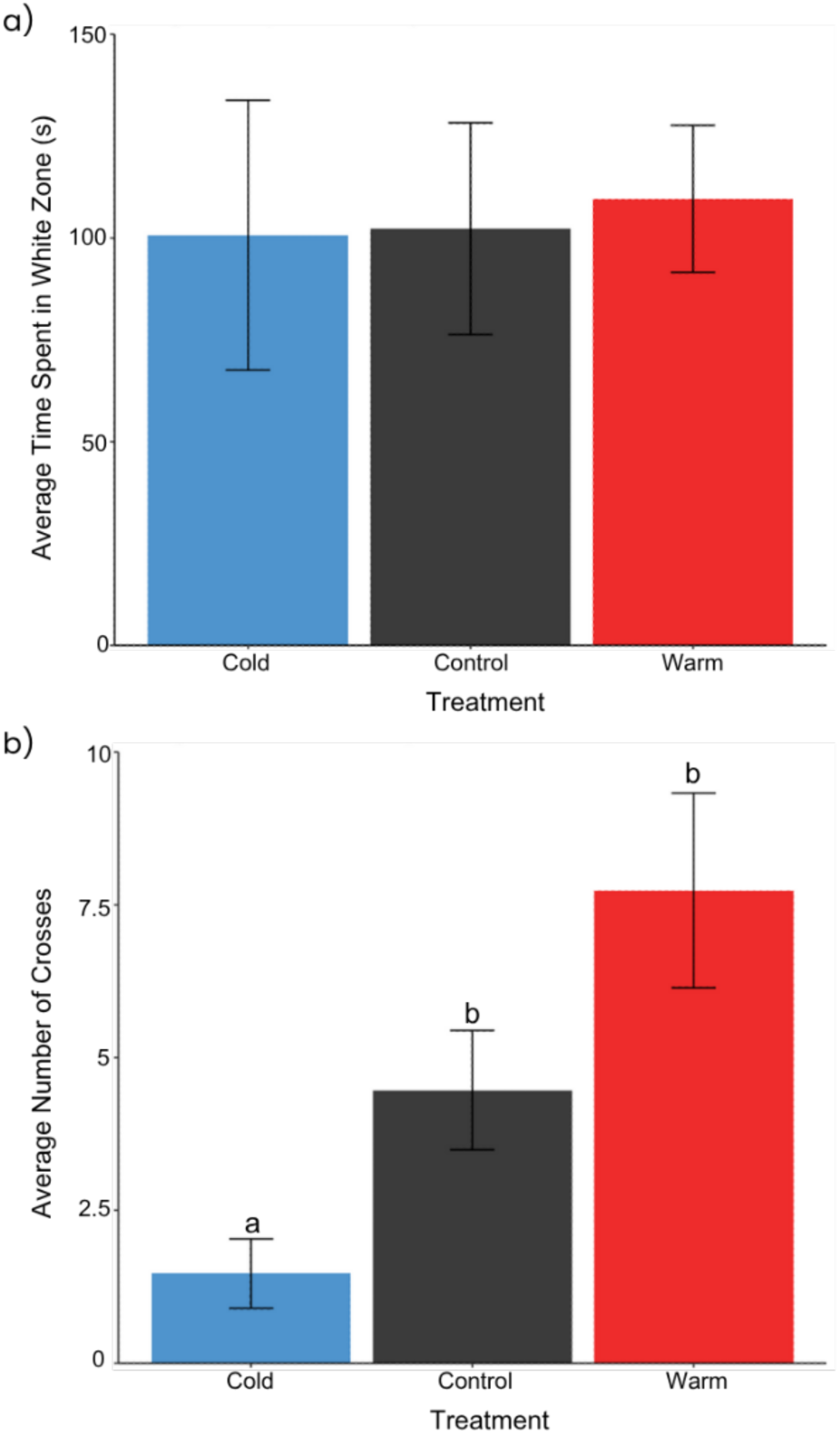
Black-White Test results for Cold (10°C), Control (15°C), and Warm (20°C) treatment groups. a) Frequency of crosses to the white zone of the tank and b) time (s) spent in the white zone of the tank (n = 15). Letters indicate significance.

In the NTT, there were no significant differences observed in the distance swam, frequency of visits to the top third of the tank, or the time spent in the top third of the tank between treatments (Kruskal-Wallis chi-squared = 1.4014, 4.182, and 5.2685; p = 0.4962, 0.1236, and 0.07177 respectively). Though not significant, in the warm treatment the average duration spent in the top third of the tank increased by about 77% and the average frequency of visits to this zone was about double that of the control (Figure 4b, c, and d). The opposite trend was observed in the cold treatment, as the average duration in the top third of the tank and the average frequency of visits to this zone were both about half that exhibited by the control (Figure 5b, c, and d). The total distance swam by the fish in both the warm and cold treatment was lower than that of the control (Figure 5a).

**Figure 5.**
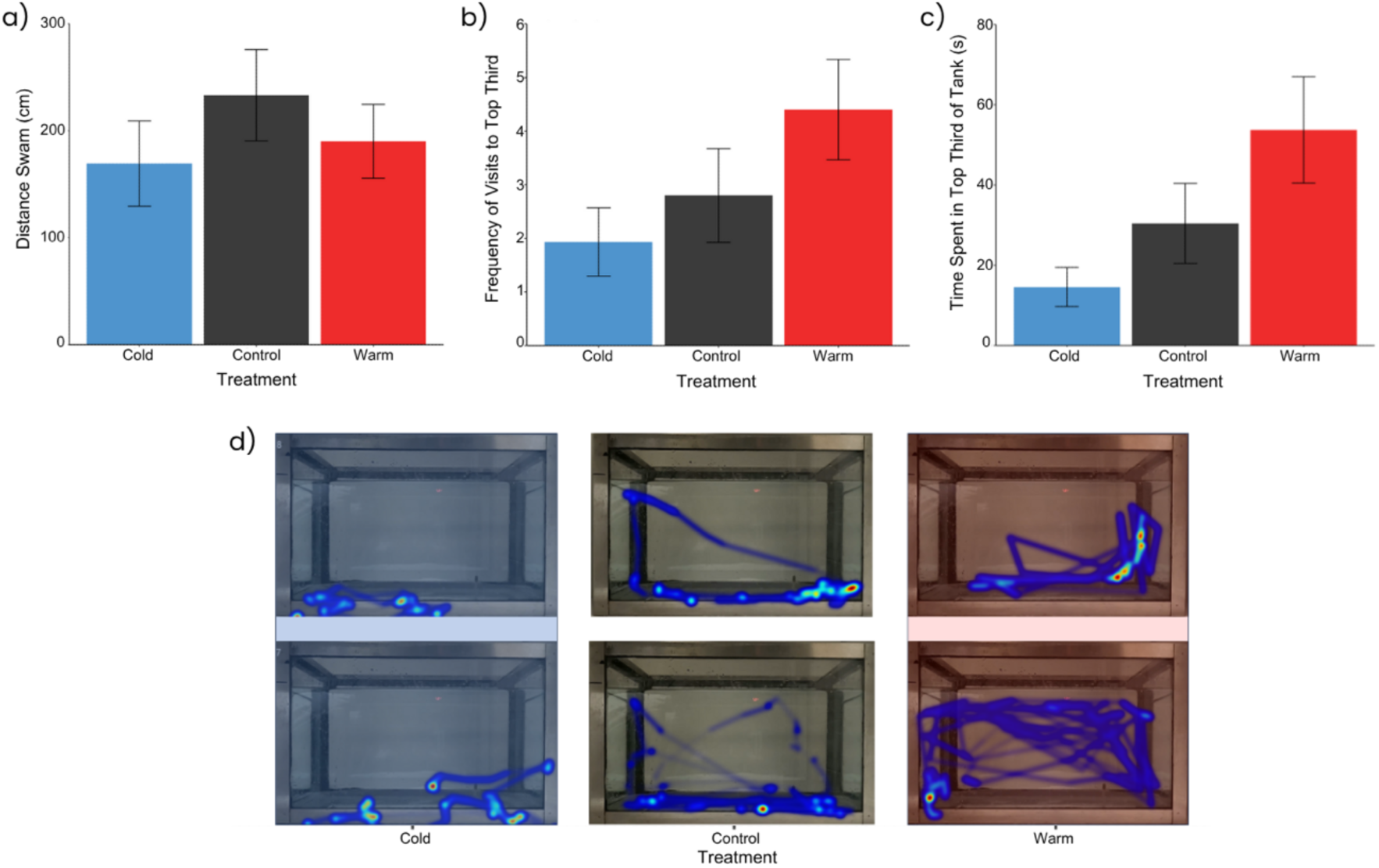
Novel Tank Test for Cold (10°C), Control (15°C), and Warm (20°C). a) Distance swam by individual (cm), b) frequency of crosses to the top third of the tank, c) time (s) spent in the top third of the tank, and d) visual representation of fish exploration in tank generated with EthoVision^®^ XT 14 where red intensity represents longer durations in area and blue represents area travelled (n = 15).

### Growth and Mortality

There was a significant increase in the change in weight in the cold treatment when compared to the control (F = 3.285; p = 0.0473), however there were no other significant differences observed between the treatments for the change in lengths, or condition factors (calculated with both exponent values; p > 0.05; see Table 1).

**Table 1.** Summary of change (before and after) in mean weight in grams, length in millimetres, and condition factor (K), calculated as 100*W*/*L*^3.04^ between treatments. Only one significant difference was found, mean weight change between cold and control temperatures, indicated by *.

| Treatment | Mean Weight Change (g) | Mean Length Change (mm) | Mean K Change |
| --- | --- | --- | --- |
| Control | 0.037 | 0.433 | 0.0057 |
| Warm | 0.072 | 0.867 | -0.0103 |
| Cold | 0.182 * | 1.133 | 0.0384 |

### Critical Thermal Maximum

There was a significant difference observed in the CT_maxs_ between all treatment temperatures (F = 169.7; p = 2e-16). CT_max_ increased with increasing acclimation temperature however the differences in CT_max_ were not proportional to the differences in acclimation temperature (Figure 6). The cold treatment had the lowest mean CT_max_ (30.95 ± 0.10 °C) followed by the control with a demonstrated a mean CT_max_ of 33.02 ± 0.08 °C. The warm treatment had the highest mean CT_max_ at 34.08 ± 0.17 °C. There were also notable behavioral differences observed between the treatment groups during the CT_max_ trials; multiple fish from the cold treatment demonstrated a jumping behavior around 20 °C, though fish were retained to their respective chambers and so CT_max_ trials were undisrupted.

**Figure 6.**
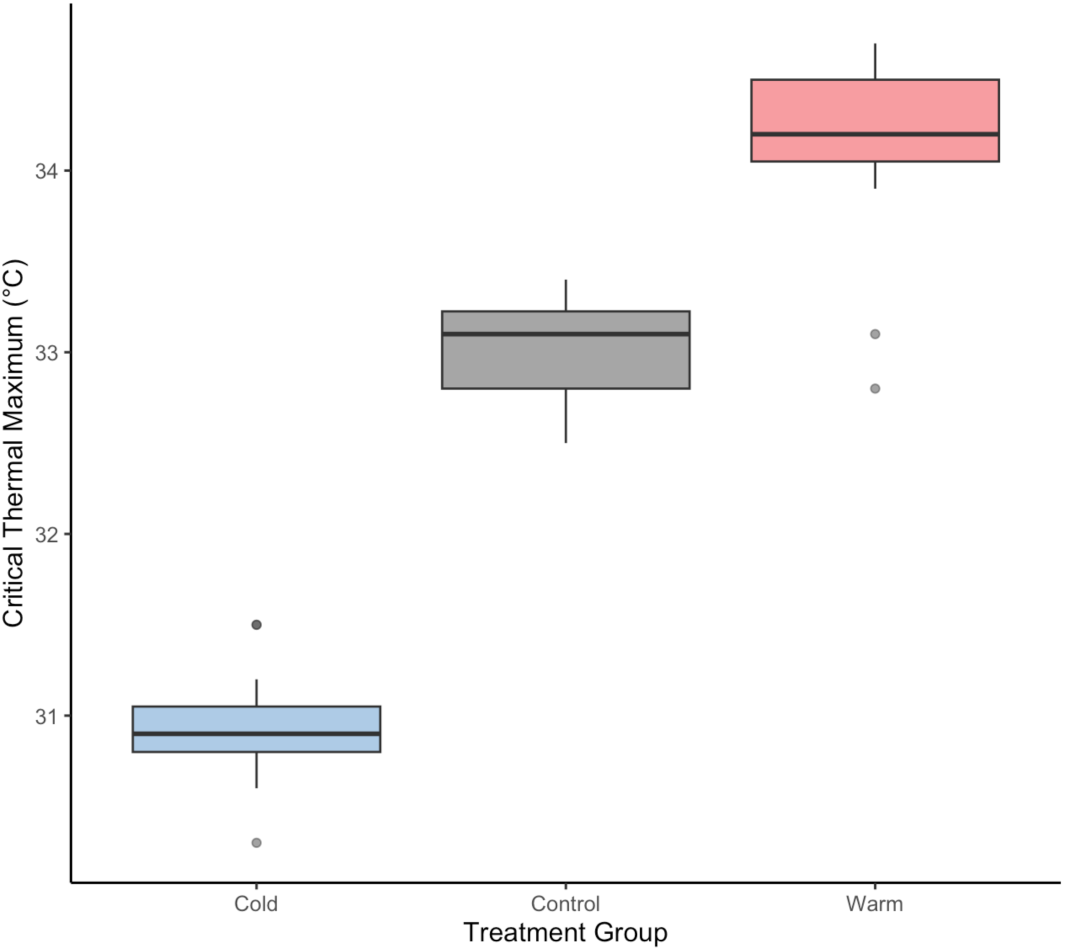
CT_max_ results for Cold (10°C), Control (15°C), and Warm (°20) treatment groups (n = 12).

### Heat Shock Protein Expression

In quantifying both HSPs 70 and 90 expression (see Figure 7) the cold acclimated fish demonstrated the least HSP expression in comparison to both the control and the warm treatment. There was no clear trend observed in regard to the HSP 90 of the warm acclimated group, though overall expression of HSP 70 was the lowest in the warm group. There were however no significant differences observed in the expression of HSPs 70 and 90 (F = 1.773 and 1.45; p = 0.219 and 0.28 respectively).

**Figure 7.**
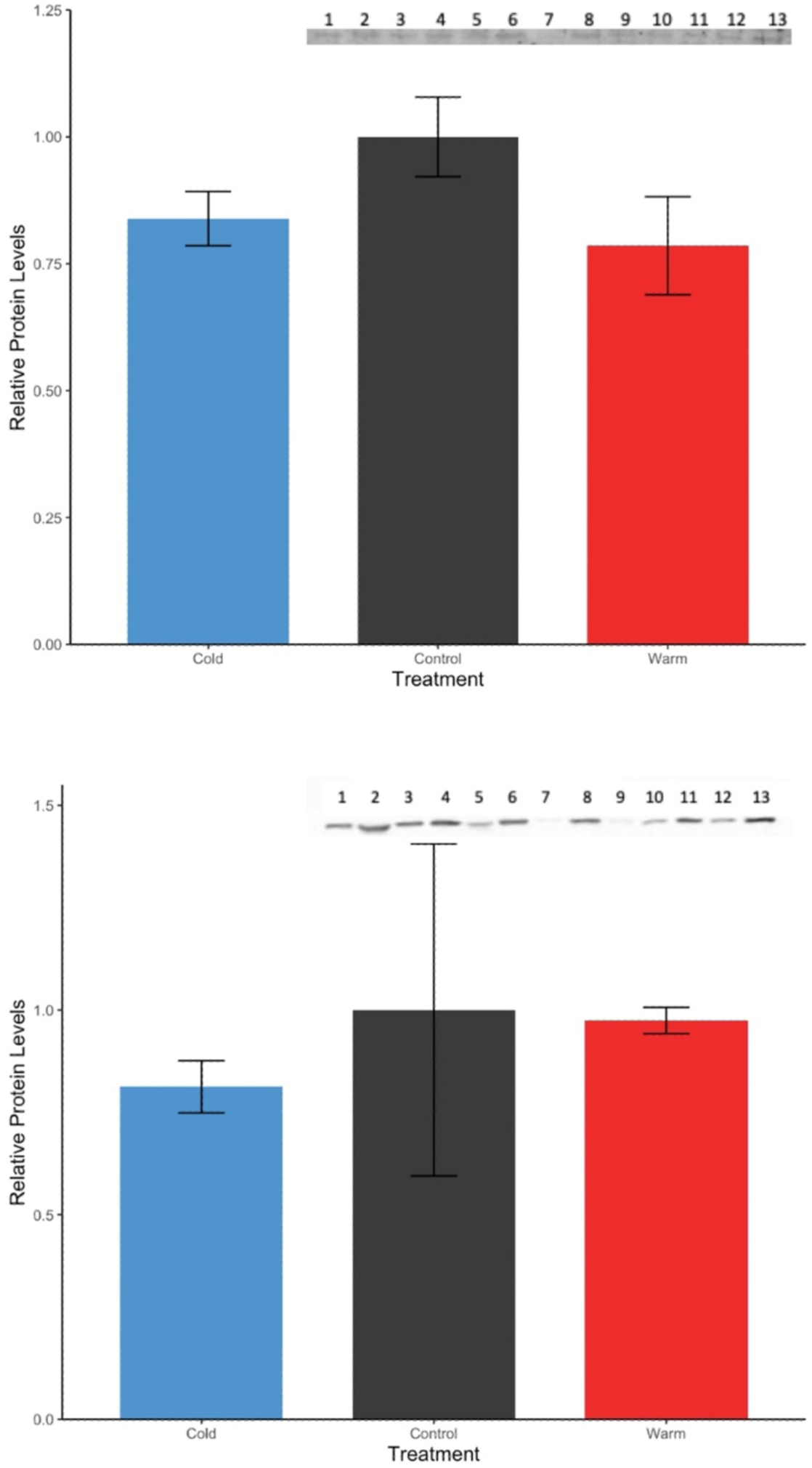
Relative protein levels for treatments Cold (10°C), Control (15°C), and Warm (20°C) for a) HSP 70 and b) HSP 90 where lanes 1-4, 5-8, and 9-13 represent cold, control, and warm expression respectively.

## DISCUSSION

As Canada warms at twice the global average and the ocean has absorbed over 90% of excess heat (3), understanding the response of fish to thermal changes is of great importance. Here we exposed threespine stickleback to conditions of 10°C, 15°C, or 20°C for 4 weeks and measured the changes in behavior, growth, CTmax, and HSP 70 and 90 content. In exploring the response of threespine stickleback across thermal ranges it is evident that behavior becomes exaggerated with rising temperature where anti-predatory behaviors decline and exploratory behaviors increase. Under warming conditions, similar to those observed over the summer season in their natural habitat, stickleback are reaching the maximums of their acclimatory potential, and while these temperatures are non-lethal, they likely cause sublethal impacts associated with the reduced performance associated suboptimal temperatures (Figure 1). These interactions likely are driving energetic trade-offs, as inferred through movement, growth, and protein content, and have potential consequences for stickleback populations considering current climate conditions and projections.

### Behavioral differences observed with temperature

Behavior is greatly important to fish on a neurological, endocrinal, and ecological level and is itself a physiological process (9). So, changes to behavior are indicators of short and long-term neurological and endocrine responses to stimuli. For example, anxiety responses indicate upregulation of corticosteroid pathways and changes to the hypothalamus-pituitary-gonadal axis (48,49).

Further, change in behavior is quite often the first (non-invasive) measurable indicator of animal response to changes in their environment (50). Because of their tight link to neurological and endocrine responses, behavior metrics may be translated through adverse outcome pathways to lower or higher levels of biological organization (51,52). Focusing on lower levels, when fish occupy and transition to habitat areas that are more susceptible and exposed (i.e., higher in the water column or brighter areas where camouflage capabilities decline) it is indicative of increased risk taking and explorative action (53). Returns to “safety” are typically characterized by anti-predatory responses such the downwards dive of fish to the bottom of a tank. These pathways are driven by the interactions of endocrinal and neurological systems, such as the movement of cortisol or the rapid release of neuropeptide-Y by neurons in the central nervous system. And likewise, looking at higher levels of biological organization, in a study done by Barolini et al. (54), changing temperatures were found to significantly impact the social and activity behaviors of the freshwater fish giant danios (*Devario aequipinnatu*). So, these behavioral indicators set the basis for breeding, population dynamics, and predatory-prey interactions, where higher anxiety and decreased sociality can starkly reduce fecundity and survival of fish (55,56). These processes establish population dynamics which in turn shape trophic relationships.

Here we observed temperature conditions significantly impact the behavior of threespine stickleback, which may have significant ecological translations. Under colder water conditions, stickleback demonstrated lower risk taking and explorative drive. Simultaneously they exhibited (anti-predatory) behaviors indicating a form of anxiety response in the new environment (57). In converse, under warming conditions, stickleback had greater risk taking and explorative drive. Interestingly these results indicate a form of behavioral differentiation occurring in response to the environmental temperature conditions, the implications of which we discuss in later sections. Notably, though, the changes we observed are significant components of common threespine stickleback behaviors like determining social hierarchies, defending territories inter- and intraspecifically, and acquiring a mate (58,59). Natural selection can directly act on these behaviors, favoring those which yield better survival, mating, and offspring outcomes (60). So, behavior and the changes we observed, are likely significant for stickleback in natural environments experiencing environmental change.

### Trade-offs associated with temperature

Trade-offs stem from a single unifying principle: energy expenditure and resource allocation is finite. Acquired resources must then be partitioned between activity, growth, reproduction, and basal metabolic activities. Due to this finitude of energy budgets, we propose that two potential trade-offs are driving our findings related to thermal acclimation.

First, we suggest that stickleback experienced an exploration-exploitation trade-off with changes in temperature. The exploration-exploitation trade-off describes the optimization of survival and resource gain considering the exploration of new environments or exploitation of known environments with established minimized risk (61). The inability to adequately “choose” inherently leads to rapid declines in resource gains and survival. Undoubtably, the lack of exploration in a natural environment and increased anxiety behavior would limit breeding success, resource assimilation, and territory defence. During the BWT, we observed that the warmed fish made a greater number of crosses to the white zone and while time spent in this zone per visit decreased. These observations could be attributed to indecisiveness and decreased problem solving, as fish are not committing to choices in crossing, which would be observed through extended periods of stay. Elevated exploration and indecisiveness may also have effects such as increasing exposure to predation and wasting energy thereby decreasing resources devoted to growth and reproduction efforts and is likely demonstrative of the exploration– exploitation trade-off (61,62).

With these changes, upper and lower trophic levels can starkly be impacted. With a prey unable to avoid predation, upper trophic levels may accidentally further limit the breeding of the proceeding generation (63). This can lead to community effects as predators are unable provision for their offspring with limited prey, thereby also detrimentally impacting their population (64). Simultaneously, the sticklebacks’ prey are allowed to thrive unchecked, with limited and ineffective predator efforts cascading these effects down the ecological trophic system. As a result, the following season may see drastic shifts in food abundance and population welfare, translatable up and down trophic levels (65).

The second trade-off we explore pertains to growth metrics. There was no significant changes found in length or condition factor between the temperature treatments; however this is not surprising as other research has demonstrated similar findings with longer timelines (27). There was however a non-significant decrease in condition factor observed in the warm treatment, though this may be attributed to energy expenditure as we generally observed that the stickleback in the warm treatment had more activity in their housing tanks. This trend in condition factor and general activity observations is supported by the growth and significant increase in weight in the cold treatment that was observed to have lower activity. This is the trade-off between activity and growth, which has been observed in lab and natural settings numerous times in multiple species, such as in the prey species roach (*Rutilus rutilus*) (66). Alternatively, the reductions in growth may be due to reduced food assimilation rates as postulated by other research with similar results (i.e., reductions in growth at warm temperatures) (67). This would imply that not all resources acquired by individuals in warmer temperatures are absorbed into the body, and increased amounts of these resources are wasted by default. Reductions in growth indicate decreased resources devoted to reproduction yielding lower progeny (68), in juveniles this may delay sexual maturity. Furthermore, the growth, weight, and condition patterns may reduce performance of fish in warmer climates during overwintering and decrease survival rates. Overall, these trade-offs represent differences in energy acquisition and expenditure dependent on temperature.

### CTmax Compression (ends of acclimatory potential)

While temperature changes in this study were sublethal, our results indicate stickleback under warming conditions are reaching the ends of their acclimatory potential. Though there was a significant difference in the CT_max_ between treatments, this difference was disproportionate to the difference in acclimation temperature. This has been observed in other species such as the brook trout (*Salvelinus fontinalis*) which demonstrated plateaus in CT_max_ shift with increasing temperatures (69). This stipulates though the acclimation temperature is not fatal, the fish is outside of its optimal temperature zone and evidently in functional decline (i.e., acclimation is limited; see Figure 1). As such performance on a cellular, physiological, and organismal level begins to decrease, with anticipated elevated apoptosis, a heat shock response, and changes to physiological blood chemistry such as increased hematocrit, plasma lactate, and glucose levels (21).These sublethal declines in performance can have substantial implications for energetic trade-offs as greater energy expenditure is required to reach similar performances, directly supporting behavioral and growth trade-offs explored above. With further warming, these trade-offs are likely to become more prominent and drive individual-population dynamics.

In the thermal limits of functionality heat shock proteins are integral in responding to and combating thermal stress. As all fish livers examined underwent critical thermal maximum (unlike most literature examining this protein content; (29)). This allowed us to explore 1) the acute-thermal stress upregulation of HSP 70 and 90 by comparison to research focused on basal states and 2) the tendency of the different temperature groups to upregulate these HSPs. To this however, there may be a degree of signal masking of HSP 70 and 90 in the warm acclimated group. Results here cannot clearly dictate whether the longer acclimation allowed for decreased HSP content and the heat shock (CT_max_) caused sudden upregulation, or the alternative.

HSP results observed here (lower expression in HSPs in both temperature extremes, with differences between HSPs) followed findings of other research. Other research on stickleback HSP expression demonstrates similar long term warm (20°C) and cold (8°C) acclimation responses with decreases in HSP 60, 70, and 90 expressions (though not significant), over time (70). We suggest from trends in the cold treatment, HSP content is reduced due to their relative lower demand at 10°C for stickleback, however upon the CT_max_ treatment the lower HSP expression is indicative of the cold acclimated group being slower to actualize needed proteins (Q_10_ effect), supported also by the lower CT_max_ values. This conclusion highlights the potential disparity between heatwave impacts in colder and warmer ambient zones. Fish inhabiting glacier fed regions or the colder regions of coast, such as the regions of Race Rocks of Canada, may face greater challenges and effects (e.g., during a heatwave), compared to those pre-exposed to warming. While the increase in HSP 70 content was more evident for the warm acclimated group in comparison to HSP 90, this may also be due to the duality of roles, in also serving as a corticosteroid carrier (71), presenting challenges in acute and long-term stress regulation. As two HSP 90 proteins are required in the heterodimer protein complex (only one HSP 70 protein), tighter homeostatic pathways exist for HSP 90, where more acute thermal stress signals are associated with HSP 70 (72), supported by our findings here. This is further supported by other studies demonstrating cortisol driven suppression of heat-stress induced HSP expression (73,74) and the higher anxiety behavioral findings of this study. There are also probable energetic trade-offs under this allostatic load driving these trends (e.g., long-term stress desensitization and energy remobilization or between HSP metabolism and fight-flight responses). While beyond the scope of our study, we recommend further investigations test multiple timepoints during exposure to different temperature conditions (with and without acute heat shock stress) to better determine the role and upregulation of HSPs during the acclimation response. Overall differences in HSP expression is directly tied to acclimation duration, heat shock, metabolism, and energetic trade-offs of fish.

## CONCLUSION

Climate change will continue to drive variability in the thermal regimes of habitats and directly impact the function and performance of animals. Here we tested the behavioral and physiological response of threespine stickleback acclimated to temperature conditions of 10°C, 15°C, 20°C for 4 weeks. We demonstrated under warming conditions that individuals exhibit increased risk-taking behaviors. Furthermore, stickleback are reaching the ends of their acclimatory capacity under current peak summer water temperatures. As a consequence, stickleback will experience trade-offs among energetically expensive processes like behavior, growth, and stress response. These trade-offs have potential consequences for stickleback populations considering current climate conditions and projections. Importantly, as higher temperature coincides with reproductive seasons, fish reproductive success, overall fecundity, and survival of populations are likely to face challenges from both the direct and indirect effects of temperature such as the energetic trade-offs (e.g., potential reductions in energy partitioning to gonads and gametes) and behavioral changes (e.g., changes to territorial defense). Stickleback are an important prey species in coastal systems, and so these effects can indirectly and directly impact lower and upper trophic levels. These trends may also be expanded to other fish species occupying a similar trophic level, driving similar effects across ecosystems as climate continues to warm. Thus, our understanding the trade-offs occurring under warming conditions will allow us to understand where populations face challenges and direct mitigation efforts accordingly.

## ACKNOWLEDGEMENTS

We thank Sean Rogers and the anonymous reviewers for their comments and suggestions which greatly improved this manuscript. Thank you also to John Post, Gary Hardiman, and Matt Vijayan for their discussions, experimental design expertise, and insight regarding this work. Thank you to the members of the Lucas, Rogers, and Jamniczky lab, Bamfield Marine Science Centre (BMSC), and the Life and Environmental Animal Care Committee (LESARC) for their help and support with animal care. Special thank you to Michael Chung, Sara Perry, Carina Lai, Enezi Khalid, and Jithine Rajeswari for their help in data collection. This work was supported by a Natural Sciences and Research Council of Canada (NSERC) Discovery Grant, NSERC Discovery Launch Supplement, and University of Calgary start-up funding to KNL. ALH was also supported by an NSERC Canada Graduate Scholarship-Masters award. The authors declare no competing or financial interests.

